# Identifying density-dependent interactions in collective cell behaviour

**DOI:** 10.1101/811257

**Authors:** Alexander P Browning, Wang Jin, Michael J Plank, Matthew J Simpson

## Abstract

Scratch assays are routinely used to study collective cell behaviour *in vitro*. Typical experimental protocols do not vary the initial density of cells, and typical mathematical modelling approaches describe cell motility and proliferation based on assumptions of linear diffusion and logistic growth. Jin *et al.* (2016) find that the behaviour of cells in scratch assays is density-dependent, and show that standard modelling approaches cannot simultaneously describe data initiated across a range of initial densities. To address this limitation, we calibrate an individual based model to scratch assay data across a large range of initial densities. Our model allows proliferation, motility, and a direction bias to depend on interactions between neighbouring cells. By considering a hierarchy of models where we systematically and sequentially remove interactions, we perform model selection analysis to identify the minimum interactions required for the model to simultaneously describe data across all initial densities. The calibrated model is able to match the experimental data across all densities using a single parameter distribution, and captures details about the spatial structure of cells. Our results provide strong evidence to suggest that motility is density-dependent in these experiments. On the other hand, we do not see the effect of crowding on proliferation in these experiments. These results are significant as they are precisely the opposite of the assumptions in standard continuum models, such as the Fisher-Kolmogorov equation and its generalisations.

## 1 Introduction

Simple two-dimensional *in vitro* experiments, such as scratch assays, are commonly used to study collective cell behaviour [1–6]. Scratch assays are conducted by placing a uniform monolayer of cells on a two-dimensional substrate and creating an artificial wound, or *scratch*, in the monolayer (figure 1*a*–*d*) [3]. Typical experimental protocols do not vary the initial density of cells between experiments and, therefore, provide no information on how the initial density affects cell migration or proliferation. In order to study potentially density-dependent cell behaviour, we consider novel scratch assay data where we deliberately vary the initial density of cells between experiments. The variation in the initial cell density in our experiments is large: the initial population in the highest density experiment is greater than the final population in the lowest density experiment.

**Figure 1:**
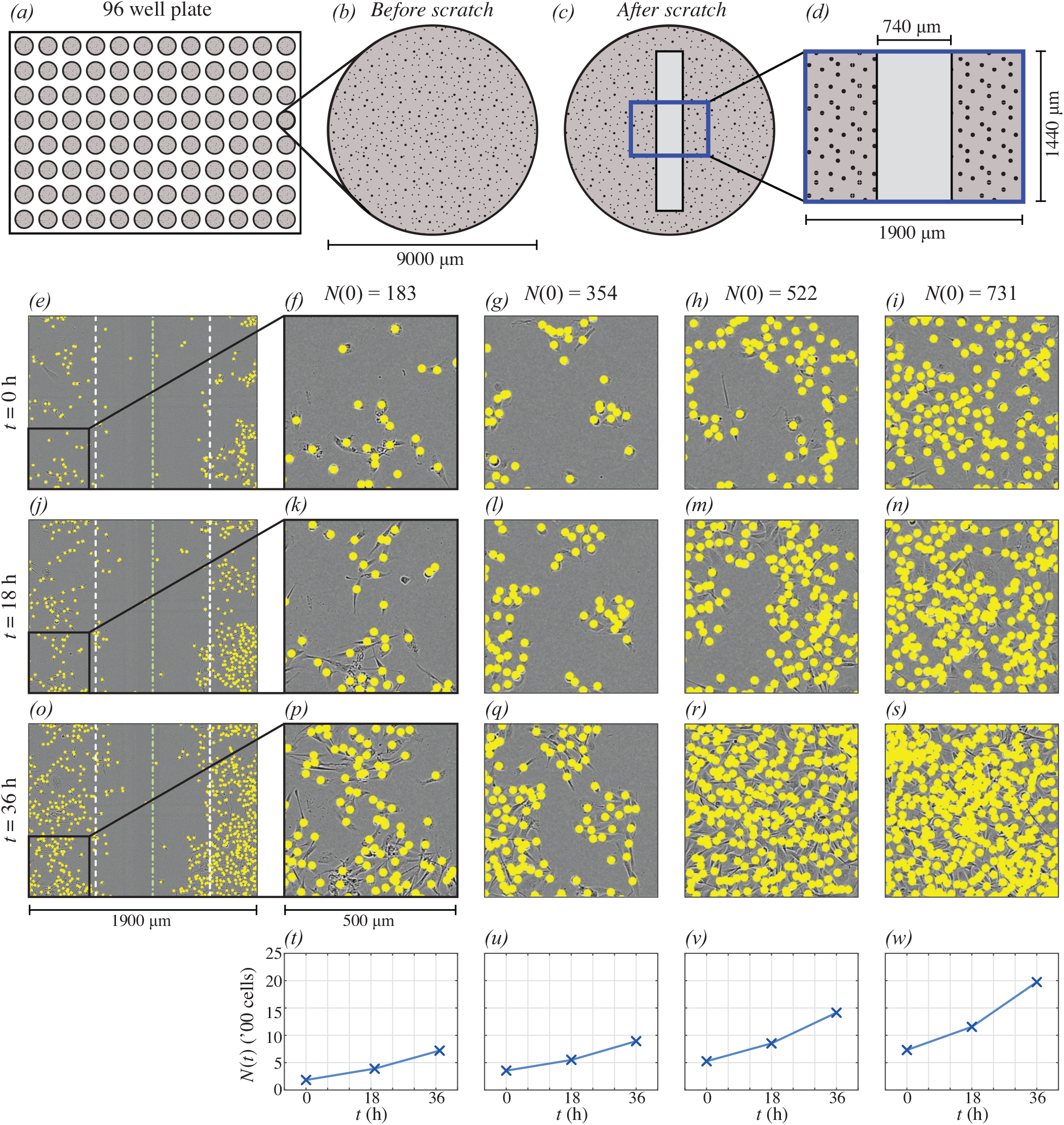
(a)–(d) Schematic of the experimental geometry. (a) 96 well plate. (b) Each assay was performed by distributing a monolayer of cells in a well of diameter 9000 μm. (c) An artificial wound (light region, not to scale) created within the monolayer of cells. (d) Field-of-view of the experimental data, which is much smaller than each well (not to scale). (e),(j),(o) Experimental data for the lowest density experiment (where *N*(0) = 183) at 0 h, 18 h and 36 h, respectively. In (e),(j) and (o) the green dash-dot line represents the approximate centre of the scratch at *t* = 0 h; and, the white dashed lines represent the approximate edge of the scratch at *t* = 0 h. Insets in (f),(k) and (p) show the lower-left region of respective images in (e),(j) and (o). The height and width of the field-of-view in the insets is 500 μm. Subsequent columns show insets for experimental data at increasing densities where *N*(0) = 354, 522 and 731. In each image, the location of each cell is indicated with a yellow marker with diameter *φ* = 24 μm (to scale). (t)–(w) Summary of experimental data in each respective column showing *N*(*t*).

Logistic growth and linear diffusion are often assumed to be the key mechanisms governing collective cell behaviour in a range of *in vitro* and *in vivo* conditions [3, 7–12]. Mean-field mathematical models that incorporate one or both of these mechanisms are routinely used to model tumour spheroids [13]; cells in living tissues [14, 15]; and simple *in vitro* experiments such as scratch [3], migration [16], and proliferation [4] assays. While calibrating these models to experimental data often leads to a good match [7], these models make the standard assumption that the parameters are independent of both initial condition and cell density. For example, the Fisher-Kolmogorov equation [17, 18]

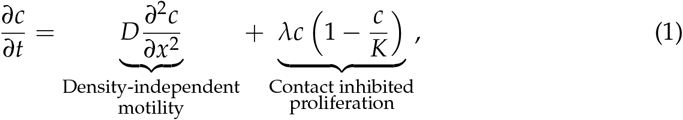

is commonly used to model scratch assay experiments [3, 19], where *c*(*x*, *t*) is the cell density. Equation (1) describes density-independent motility, characterised by a constant diffusivity *D*; and density-dependent proliferation, characterised by a constant proliferation rate *λ* and a constant carrying capacity *K*. Jin *et al.* [3] find that calibrating the solution of equation (1) to scratch assay data yields vastly different estimates of *D* for each initial condition considered. In contrast to equation (1), some studies assume that cell motility is density-dependent. However, the way in which this is modelled is inconsistent. For example, Cai *et al.* [20, 21] model motility with a non-linear diffusivity term that decreases with density to simulate crowding. In direct contrast, many other studies model motility using a non-linear diffusivity term that increases with density to simulate contact stimulation [22–24]. These studies all calibrate their respective models to experimental data with a single initial density [20–24]. In contrast, the approach that we take here is biologically significant since we identify the nature of density-dependent interactions using experimental data initiated with a range of initial cell densities.

In this work we describe the cell behaviour with a lattice-free individual based model (IBM) [4, 25, 26]. The IBM represents cells as *agents* that take locations in continuous space, and so we can specify the initial agent locations in the model to precisely match the initial cell locations in the experiments. This choice also allows the model to capture local details—such as spatial structure and clustering—which are neglected by standard continuum modelling approaches [3, 10]. The agents in the IBM proliferate and move, the rates of which we assume depend explicitly on interactions between neighbouring agents. We quantify these interactions with kernels that depend on the distance between pairs of cells. Directional bias is also incorporated so that agents are more likely to move either away from, or towards, regions of high density [21, 25]. A key advantage of the IBM is its flexibility: it is trivial to add and remove mechanisms, which we do to study the interactions required for the model to simultaneously match all experiments. Finally, the IBM is stochastic and so naturally describes the variation between experiments.

We aim to identify the nature of interactions which enable the model to simultaneously describe experimental data across a wide range of initial cell densities. This study is the first time scratch assay data initiated across a range of initial cell densities has been calibrated to an IBM. We take a Bayesian approach to parameter estimation [4, 27–29], and identify interactions using model selection [27]. We always force the model to simultaneously match data from all nine experiments. The mathematical model is always initiated using the initial configuration of cells in each experiment, and we compare simulated and experimental data at 18 h and 36 h, the latter which corresponds to the duration of the experiment. The calibrated model is able to replicate the experimental data, and we find evidence to suggest that motility is an increasing function of density, which is contrary to both the common mathematical assumption of linear diffusion and work which assumes motility decreases with density [20, 21]. Experimentation with summary statistics confirms the importance of spatial structure, which is neglected by standard modelling and model calibration approaches.

## 2 Materials and methods

### 2.1 Experimental methods

Our experimental model of cell migration and proliferation comprises a series of scratch assays using PC-3 prostate cancer cells [30]. We deliberately vary the initial number of cells in each experiment by seeding approximately 8000, 10000 and 12000 cells in a 9000 μm diameter well within a 96-well plate (figure 1*a,b*). Cells are grown overnight to create a spatially uniform monolayer before a scratch is created (figure 1*c*). Images of the central 1440 × 1900 μm of each well are captured over a period of 48 hours after the monolayer is scratched (figure 1*d*). Full details of the experimental methods are provided in [3].

ImageJ [31] is used to determine the approximate coordinates of individual cells in each image, this data is given in the supporting material. We exclude the first 12 hours of experimental data from our analysis [4] to ensure that sufficient time has passed so that the cells are migrating and proliferating after the scratch has been made. We then record experimental images and we treat this as the beginning of the experiment, *t* = 0 h. The variability in initial cell number is high: despite an initial seeding density of approximately 8000–12000 cells per well, which corresponds to expected initial number of cells within the field-of-view of 344–516, we find that the initial number of cells within the field-of-view at *t* = 0 h ranges from 183 to 731 (figure 1*f-i*). This variation is also high between experiments of the same seeding density [32], due to the fact that our field-of-view is relatively small so that fluctuations about the expected values are relatively large. We demonstrate this variation in figure 1*e–s*.

### 2.2 Mathematical model

We use a lattice-free individual based model (IBM) [4, 25] which we simulate with the Gillespie algorithm [33]. The model includes density-dependent proliferation and movement, but does not consider death, which is not observed in the experiments. To be consistent with previous experimental observations [21], the model incorporates a bias mechanism so that cells both move, and disperse daughter cells during proliferation, in a direction either towards, or away from, crowded regions. Density dependence is incorporated into the model through cell-to-cell interactions, so that behaviour is dependant upon *local crowding*.

The field-of-view of the experimental data is rectangular, with dimensions 1440 × 1900 μm (figure 1*d*), and we replicate this by using the same geometry in the model. As the well in the tissue culture plate is much larger than this field-of-view, we apply periodic boundary conditions [4] (indicated in blue in figure 1*c,d*). Cells are modelled as *agents* that have a point location but no physical size. In our previous work we find that, on average, these PC-3 prostate cancer cells have an area that corresponds to a disc of diameter *φ* = 24 μm [4]. The interaction mechanisms we model are not based on volume exclusion or hard sphere interactions [26], but rather depend on agent separation in such a way that configurations wherein two agent centres are very close are unlikely. We denote the agent locations **x**_*n*_ = (*x_n_*, *y_n_*), *n* ∈ {1, …, *N*(*t*)}, where *N*(*t*) denotes the number of agents in the simulation. We specify the initial agent locations in each simulation to match the experimental images at *t* = 0 h.

#### Directional bias

We quantify crowding by placing a *bias kernel* at the location of each agent to form a *crowding surface*, *B*(**x**), as shown in figure 2*c,d* for the configuration of cells in figure 2*a,b*. Mathematically, this is given by

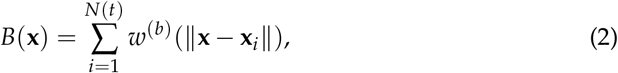

and describes a measure of local crowding at **x**, where *w*^(*b*)^(*r*) is the bias kernel. The contributions of each agent to *B*(**x**) depend on the distance between **x**and the location of the *i*th agent, **x**_*i*_, given by *r* = ‖**x** – **x**_*i*_‖. There are many possible choices of kernel [34], however we find that the standard choice of Gaussian leads to a good match with experimental data [4]. In this study, we choose *w*^(*b*)^(*r*) to be a Gaussian [35] of spread *σ* with an extremum of *γ*_*b*_ so that

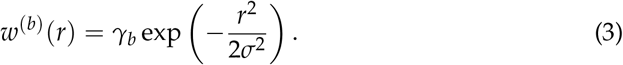

**Figure 2:**
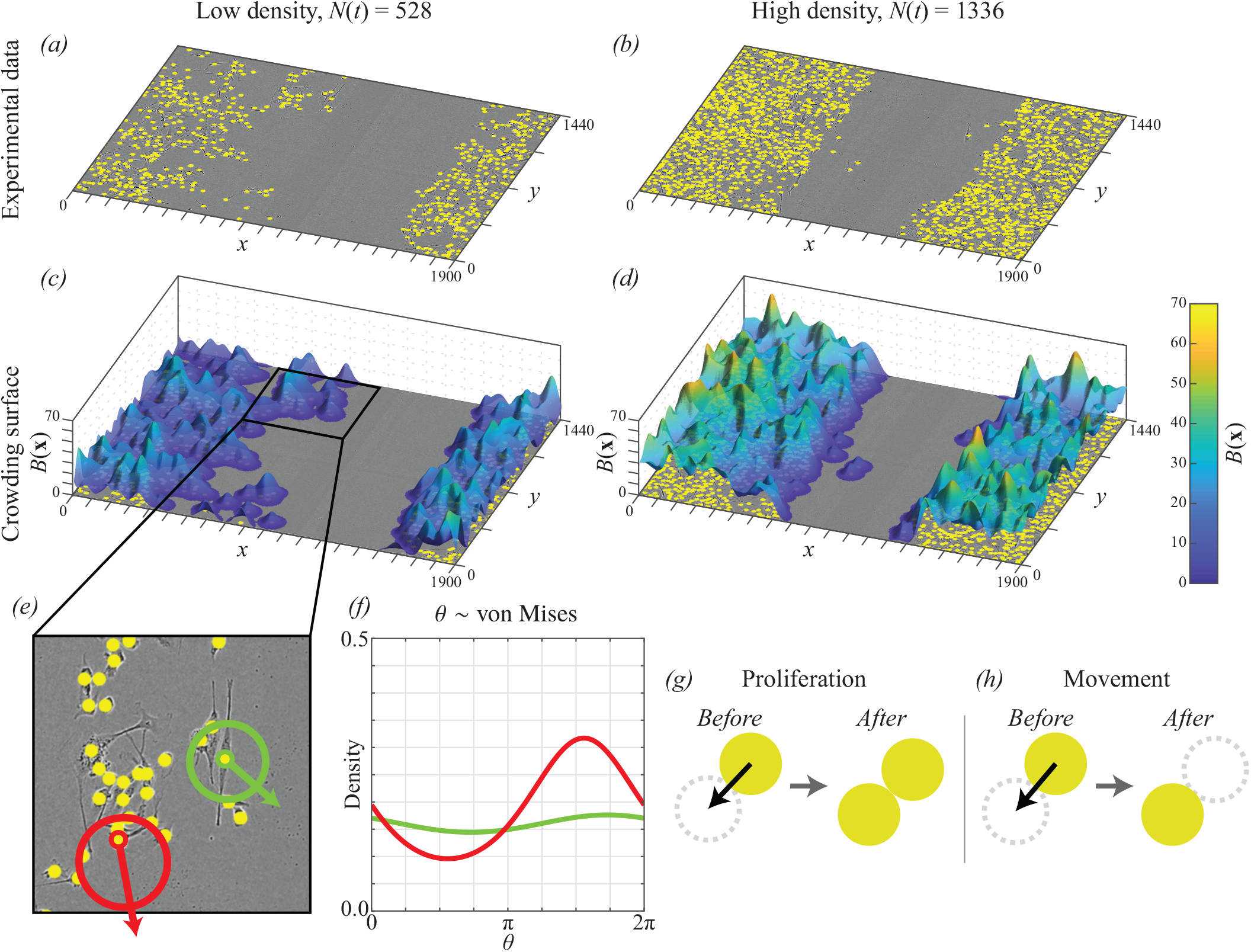
(a)–(b) Experimental data tat (a) low density; and, (b) high density. The location of each cell is indicated with a yellow marker of diameter *φ* = 24 μm (to scale). (c)–(d) Example crowding surface, where agent locations are taken from experimental data in (a) and (b), respectively. (e) Two-dimension inset of experimental image in (c), showing the bias distribution for two agents in radial coordinates centred at each agent. The off-centredness of each circle therefore represents the strength of the bias, which is stronger for the red cell than the green cell. (f) The bias distributions in (e) shown in Euclidean coordinates for clarity. (g)–(h) Schematic of cell division (proliferation) and movement events, respectively, where the black arrow indicates the sampled direction of each cell. When an agent proliferates, the daughter cell is placed a distance of *φ* from the mother cell. When an agent moves, the agent is moved a distance of *φ*.

For computational efficiency, we truncate the kernel to zero for *r* ≥ 3*σ* [35]. This truncation means that agents separated by a distance of more than 3*σ* do not interact. Therefore, *B*(**x**) is a measure of *local crowding*.

For *γ*_*b*_ > 0, agents prefer to move and disperse daughter agents in the direction of steepest descent on the crowding surface, which corresponds to regions of lower density (setting *γ*_*b*_ < 0 has the opposite effect). This preference depends on the steepness, so that agents close to highly crowded regions are more likely to move and disperse daughter agents in their preferred direction, demonstrated in figure 2*e,f*, where the red agent has a stronger bias strength than the green agent. To do this, we define the bias vector of agent *n* as

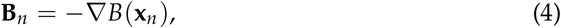

which gives the magnitude and direction of steepest descent. Therefore, **B**_*n*_ is a simple measure of local spatial structure at the location of agent *n*. The movement and proliferation directions are then sampled from the von Mises distribution [36]

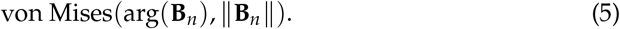

The expected and most likely direction is, therefore, arg(**B**_*n*_). The direction distribution becomes increasingly concentrated around arg(**B**_*n*_) as ‖**B**_*n*_‖ becomes large, and approaches a uniform distribution on [0, 2*π*) as ‖**B**_*n*_‖ → 0.

We illustrate the directional bias mechanism in figure 2*c*-*f*. The crowding surface is constructed by placing a Gaussian kernel at the location of each agent (figure 2*c,d*). In figure 2*e* we show the bias distribution and preferred direction for an agent in a low (green) and high (red) density region. For each agent the arrow shows the preferred direction with the corresponding von Mises distribution plotted in radial coordinates centred at the location of each agent. In figure 2*f* we show these distributions are shown as a function of the angle, *θ* ∈ [0, 2*π*), for clarity.

#### Proliferation and movement

Cell division (proliferation) and movement events occur according to a Poisson process [37] with density-dependent rates *P*_*n*_ ≥ 0 and *M*_*n*_ ≥ 0, respectively. These rates comprise constant intrinsic rates *p* > 0 and *m* > 0, that are modified by interactions with neighbouring agents. These interactions result in a local density dependence, so that agents in high density regions are able to behave differently to solitary agents, or agents in low density regions [25].

We quantify these interactions using kernels, *w*^(·)^(*r*), that depend on the separation distance, *r* ≥ 0, between an agent and its neighbours, such that

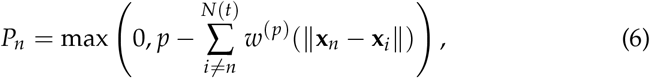

and

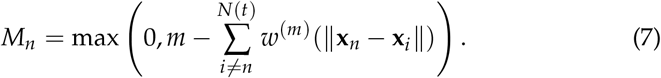

Again, we choose the kernels to be Gaussian [35], with spread *σ*, so that

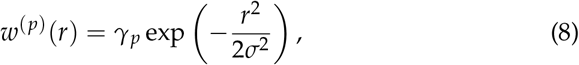

and

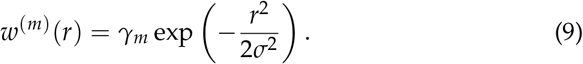

Here, *γ*_*p*_ and *γ*_*m*_ are the extrema of the proliferation and movement kernels, respectively. Setting *γ*_*m*_ < 0 (or *γ*_*p*_ < 0) means that crowding increases motility (or proliferation); setting *γ*_*m*_ > 0 (or *γ*_*p*_ > 0) means that crowding decreases motility (or proliferation); and, setting *γ*_*m*_ = 0 (or *γ*_*p*_ = 0) means that motility (or proliferation) is independent of local density. Again, we truncate the kernels to zero for *r* ≥ 3*σ* [35].

When an agent at **x**_*n*_ proliferates, the daughter agent is dispersed a distance *φ* (approximately one cell diameter) from **x**_*n*_, with the direction sampled from the bias distribution for that agent (figure 2*e,f*). This is demonstrated in figure 2*g*. When an agent at **x**_*n*_ moves, it is moved to a location of distance *φ* from **x**_*n*_, with the direction sampled from the bias distribution for that agent (figure 2*e,f*). This is demonstrated in figure 2*h*.

### 2.3 Summary statistics

To match model simulations to the experimental data, we record the locations of agents at both *t* = 18 h and *t* = 36 h. We denote the experimental data at both time points from experiment *i* ∈ {1, …, 9} as 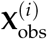, and simulation data from experiment *i* as 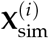. In this section, we detail how we summarise the high dimensional data **X** into lower dimensional summary statistics. This allows us to define a distance function, *d*(**X**_obs_, **X**_sim_), that represents the distance between experimental and simulation data.

We aim to capture three key pieces of information in the experiments: (1) the population size; (2) the spatial structure; and, (3) the density profile. The first two pieces of information are related to the first two spatial moments [35], and the last piece of information relates to the wound closure, total population and the spatial distribution of cells. The first spatial moment, the average density, is the number of agents in the population, *N*(*t*). The second spatial moment describes the spatial distribution of agents, often characterised by a pair correlation function [2, 25, 35]. In summary, the pair correlation function describes the density of pairs of agents separated by a distance *r*, relative to the expected density of pairs if the population were uniformly distributed [2]. Since the data is discrete, we define the pair correlation, 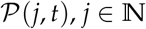, which describes the relative number of pairs separated by a distances ranging from (*j* – 1)Δ*r* < *r* < *j*Δ*r*, given by

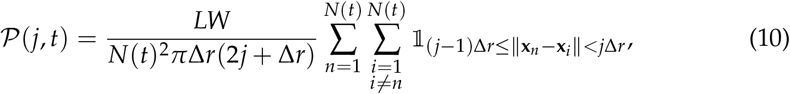

where *L* and *W* are length and width, respectively, of the region and 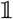 is the indicator function. In this study, we choose Δ*r* = 5 μm, and consider the pair correlation up to a distance of 100 μm such that *j* ≤ 20. Smaller values of Δ*r* lead to a noisier pair correlation function, and larger values of Δ*r* smooth the pair correlation, potentially hiding information [38].

In a scratch assay the central region of the experimental field-of-view is initially devoid of agents (figure 3*a*). To account for this, we calculate pair correlation functions for sub-region of width 400 μm in the far-left, and far-right, of the domain (figure 3*a*, indicated in red) denoted 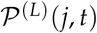 and 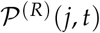, respectively. The width of this region is chosen so it does not overlap with the region devoid of cells in the experimental images. We apply periodic boundary conditions on these sub-regions, so that the separation of a pair of agents is the smallest possible distance accounting for the periodic boundary conditions. The pair correlation function that summarises the entire experiment is 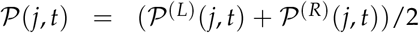 (figure 3*b*). Results in *figure 3b* also confirm our assumption that a typical cell diameter is approximately 24 μm [4]. Results in the supporting material (figures S8 and S10) show the pair correlation function for all experimental images, where clustering at short distances is observed for earlier time data, for lower cell densities.

**Figure 3:**
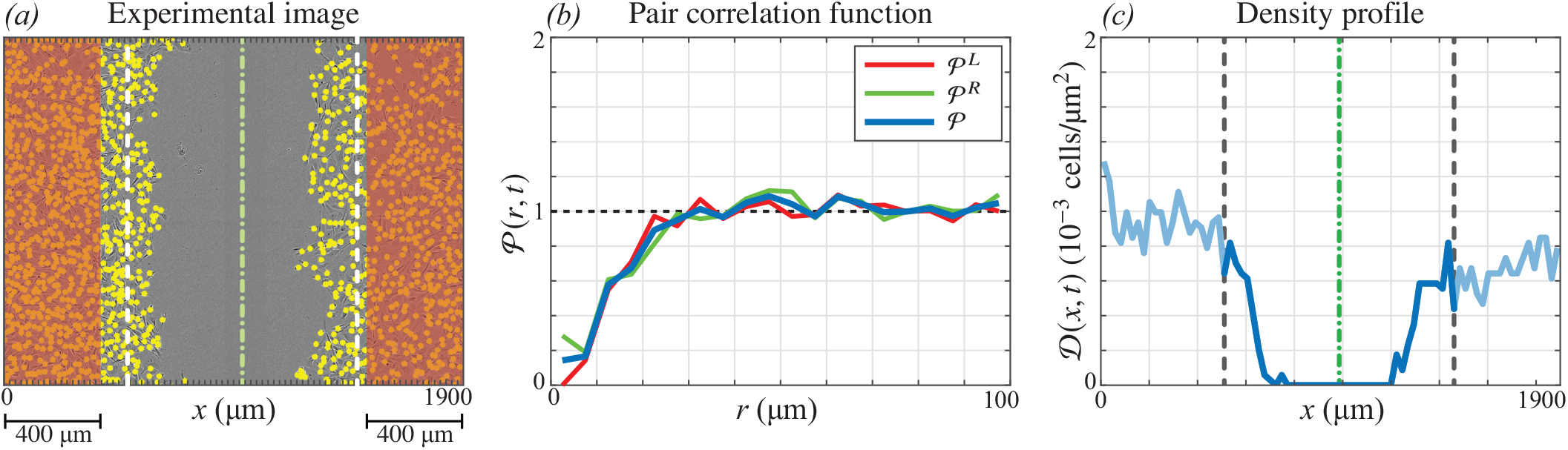
(a) Experimental image. The location of each cell is indicated with a yellow marker of diameter *φ* = 24 μm (to scale). (b) Pair correlation function calculated from the distribution in cells in (a). 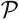 is the average of 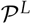 and 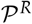, calculated using the agents in the 400 μm to the far-left and far-right (red region) of the experimental data in (a), respectively. Therefore, 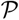 contains information about the spatial distribution of cells, but not about the scratched region. Pair correlation functions for all experimental images are provided in the supporting material (figures S8 and S10). (c) The density profile, calculated by counting the number of cells in subregions of width 1900/80 μm and dividing by the area of each subregion. The sub-regions are indicated as the axis ticks in (a). Only the central 41 bins are used to compare experimental and simulated data. Green dash-dot lines in (a) and (c) indicate the approximate centre of the scratch at *t* = 0 h and dashed lines indicated the approximate boundary of this region at *t* = 0 h.

The final piece of information, the density profile, describes the wound closure, total population and spatial structure. We subdivide the field-of-view in figure 3*a* into 80 vertical sub-regions, each of width Δ*x* = 1900/80 = 23.75 μm. We define the density profile 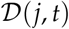 to be the number of agents with an *x*-coordinate between (*j* – 1)Δ*x* and *j*Δ*x*, divided by the area of the sub-region, giving the density. This density profile is shown in figure 3 *c*. To avoid capturing excessive noise in our measurement of wound closure, we do not include the entire density profile in the distance metric. Rather, we manually approximate the *x*-coordinate of the centre of the scratch at *t* = 0 h for each experiment, denoting 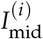 as the bin index of the centre the scratch in experiment *i*. We include the central 41-subregions which, in effect, surround the initially scratched region of each experiment. This region is indicated in figure 3*c* and avoids the fluctuations in density outside this region.

The distance metric, *d*(**X**_obs_, **X**_sim_), is given by

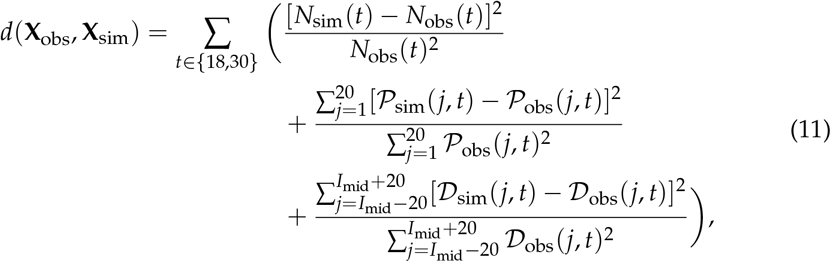

and includes information from all three summary statistics, at *t* = 18 h and *t* = 30 h. Therefore, *d*(**X**_obs_, **X**_sim_) is the relative square error of the simulation from the experiment. For 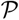 and 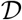, the contributions to *d*(**X**_obs_, **X**_sim_) approximate the relative square error in the integral of each summary statistic, given the spatial discretisation we have applied to each.

### 2.4 Approximate Bayesian computation and model selection

We consider a hierarchy of models. The full model, which we denote as Model 1, contains the five unknown parameters *θ*_1_ = (*m*, *p*, *γ_m_*, *γ_p_*, *γ*_*b*_). Models 2 to 5 are subsets of the full model, where we progressively restrict various combinations of the interaction strength parameters *γ_m_*, *γ_p_* and *γ*_*b*_ to be zero, effectively removing them from the model. We summarise these five models in table 1, where we denote *θ*_*k*_ as the unknown parameter combination for Model *k*.

**Table 1:**
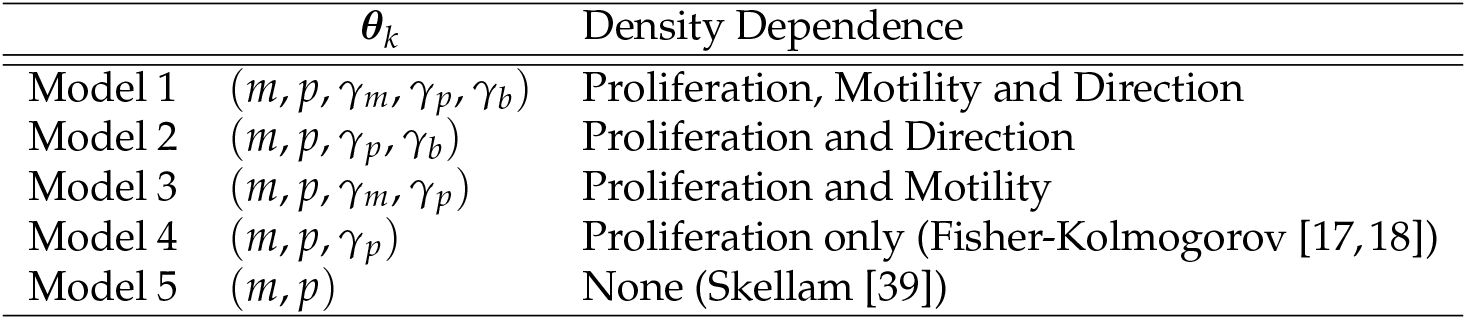
The hierarchy of models considered. The full model (Model 1) contains a parameter governing the: motility rate, *m*; proliferation rate, *p*; motility interaction strength, *γ*_*m*_; proliferation interaction strength, *γ*_*p*_; and, directional bias strength, *γ*_*b*_. In subsequent models, we restrict various combinations of the parameters to zero, effectively removing them from the model.

We treat the unknown parameters in each model as a random variable, *θ*. In the absence of experimental observations, our knowledge of *θ* is characterised by specified prior distributions. When included in the model, the priors were chosen to be independent and are as follows: *π*(*m*) = *U*(0, 10)/h; *π*(*p*) = *U*(0.02, 0.05)/h; *π*(*γ_m_*) = *U*(−2, 2)/h; *π*(*γ_p_*) = *U*(0, 0.02)/h; and *π*(*γ*_*b*_) = *U*(0, 100) μm. In this context, *π*(·) represents a probability distribution. In the supporting material, we show that widening these priors has negligible effect on the results. Initially, we also treat *σ* as an unknown parameter where *π*(*σ*) = *U*(2, 30) μm. This initial analysis provides strong evidence for the value of *σ*, so we set *σ* = *φ*/2 = 12 μm to decrease the dimensionality of the parameter space. In the supporting material, we also investigate *σ* = *φ* = 24 μm, since this is a natural choice in a lattice-based framework where the migration distance and dispersal distance are also the same as the average agent diameter. We apply approximate Bayesian computation (ABC) [4, 15, 27, 29] to update our knowledge of the parameters using experimental observations, 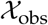, from all nine experiments, to produce posterior distributions, 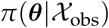. Since this model is known to be computationally expensive [4] and we have a high-dimensional parameter space, we apply an ABC method based on sequential Monte-Carlo (SMC) [27, 29, 40].

In this study, we aim to find parameter combinations that simultaneously match all nine experimental data sets, such that 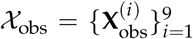. For each prior sample in the ABC rejection algorithm we simulate a model realisation using each experimental initial condition, to obtain 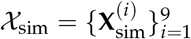. We then compare observed data, 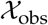, to simulated data, 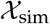, using the discrepancy measure

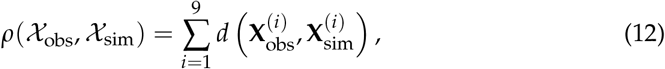

where *d*(·, ·) is given in equation (11). In ABC techniques, we accept a proposal as a posterior sample if 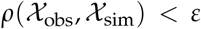 for some threshold *ε*. As *d*(·, ·) ≥ 0, the sum in Equation (12) is non-decreasing in *i*. We therefore implement early rejection [41] by sequentially producing model realisations for *i* ∈ {1, …, 9}. If, at any time, the partial sum up to a value *i* exceeds the threshold *ε*, we immediately reject the sample. In practice, this saves considerable computation time by reducing the number of times the model must be simulated using high-density initial conditions.

The principle behind ABC SMC is to propagate a series of prior samples, called *particles*, through a sequence of distributions 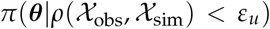, *u* = {1, …, *U*} [27, 29, 40]. The thresholds *ε_u_* satisfy *ε*_*u*_ > *ε*_*u*+1_, so that the distribution gradually evolves to the target distribution 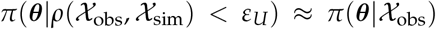. To obtain a sequence of thresholds, and an estimate of the smallest discrepancy possible in all models, we first perform a pilot run using ABC rejection [4, 29] with Model 1 (supporting material, section 1.1). From 100,000 prior samples, this provides an estimate of the probabilities 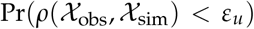, given *θ* is simulated from the prior. We choose the sequence 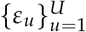 by examining a quantile plot (supporting material, section 3). We choose *ε_U_* to corresponds to an acceptance rate of approximately 1% under ABC rejection. The sequence of discrepancies, and details of the ABC rejection and SMC algorithms are given in the supporting material (sections 1 and 3).

We follow the ABC SMC algorithm of Toni *et al.* [27] to perform parameter inference and model selection. Under this algorithm, we place a prior distribution on the model index, *π*(*M_k_*), which we choose to be a discrete uniform distribution so that each model is equiprobable. ABC SMC is then used to estimate the posterior probability of each model, 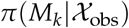. We detail this algorithm in the supporting material (section 1.2). A key feature of this technique is to implicitly penalise models with a higher number of parameters. We compare models by computing the Bayes factor, 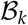 [42], which describe the *evidence* in favour of Model *k* over the full model, Model 1. As a uniform prior is placed on the model index, the Bayes factor is given by

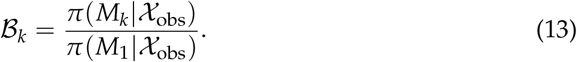

Here, 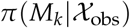 denotes the marginal posterior density of *M_k_* (Model *k*). A value 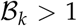 indicates evidence in favour of Model *k* compared to the full model, and viceversa for 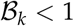. The Bayes factor is therefore simply the ratio of the posterior density for Models *k* and 1, and provides evidence to compare models in a similar way to that used in frequentist hypothesis testing.

## 3 Results and Discussion

Common mean-field models, such as the Fisher-Kolmogorov equation [18] and its generalisations, are not able to simultaneously describe collective cell behaviour in scratch assay experiments across a range of initial densities [3]. This suggests density-dependent behaviour in these experiments. Our model allows interactions between cells to affect proliferation, movement and direction. To identify the importance of each of these interactions, we simultaneously calibrate our model to nine scratch assay experiments which we initiate across a wide range of initial densities. We always initiate the IBM using the initial configuration of cells in the experiments and perform inference using data at an intermediate time point, *t* = 18 h, and at the conclusion of the experiment, *t* = 36 h. In a set of preliminary results (not shown) we only included data at the last time point, *t* = 36 h, and the inclusion of the intermediate time point made negligible difference to the results. Therefore, we do not expect the results to change significantly should more than two time points be considered.

Our first result is to identify the distance over which these interactions occur. We quantify interactions using Gaussian kernels dependent on the distance between pairs of agents [35], and characterised by a spread parameter *σ* (equations (3), (8) and (9)). The interaction between a pair of agents separated by more than approximately 3*σ* is, therefore, negligible. We expect *σ* to be of the same order of magnitude as *φ* = 24 μm, which is the approximate cell diameter [4]. We perform ABC rejection where *σ* is sampled from the prior *U*(2, 30) (supporting material, section 1.1). These results suggest that *σ* ≈ *φ*/2 = 12 μm, and we fix this for the rest of the study to reduce the number of unknown parameters. This result suggests that interactions between cells occurs over a relatively short distance, since the model predicts interactions between cells separated by more than 3*σ* = 36 μm is negligible.

One of the most important aspects of the lattice-free IBM is its ability to describe, in fine detail, the spatial structure of cells in the experiments, which we quantify using the pair correlation function. In contrast, mean-field models consider only average properties of the cell population [19] and lattice-based methods [26, 28] are not able to precisely capture the initial agent configuration from the experiments. Lattice-based methods also, by definition, constrain the separation of agents to take discrete values, and typically agents in these models cannot lie closer than one cell diameter. The pair correlation describes the probability of finding pairs of agents separated by each distance, and hence can provide information about the effect of interactions on the dynamics. To show this, we repeat ABC rejection but exclude the pair correlation function from the distance metric (supporting material, section 2.3). These results show that the posterior distributions change significantly in this case, verifying that the pair-correlation function contains a significant amount of information about these interactions.

To quantitatively determine the importance of each interaction, we consider a hierarchy of models where we successively set interaction strength parameters (*γ_m_*, *γ_p_* and *γ*_*b*_) to zero to remove the corresponding interaction from the model. We use the model selection algorithm of Toni *et al.* [27], and compare the evidence in favour of each model over the full model (Model 1) using Bayes factors [27, 42]. We show the posterior density for each model in figure 4*a*, and summarise the Bayes factors and evidence in table 2. Overall, we find that Model 1 has the highest posterior density (figure 4*a*). We find positive evidence in favour of Model 1 over Model 2 (where *γ_m_* = 0 and so motility is density-independent); and weak evidence in favour of Model 1 over Model 3 (where *γ*_*b*_ = 0 and so there is no directional bias). Importantly, we find that Models 4 and 5, where *γ_m_*, *γ*_*b*_ = 0 and *γ_m_*, *γ_p_*, *γ*_*b*_ = 0, respectively, cannot match the experimental data 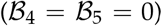. Contrary to assumptions that are commonly made in models such as the Fisher-Kolmogorov equation, these results provide evidence to suggest that motility is density-dependent, as either a density dependent movement rate must be included (Models 1 and 3) or a directional bias (Models 1 and 2).

**Table 2:**
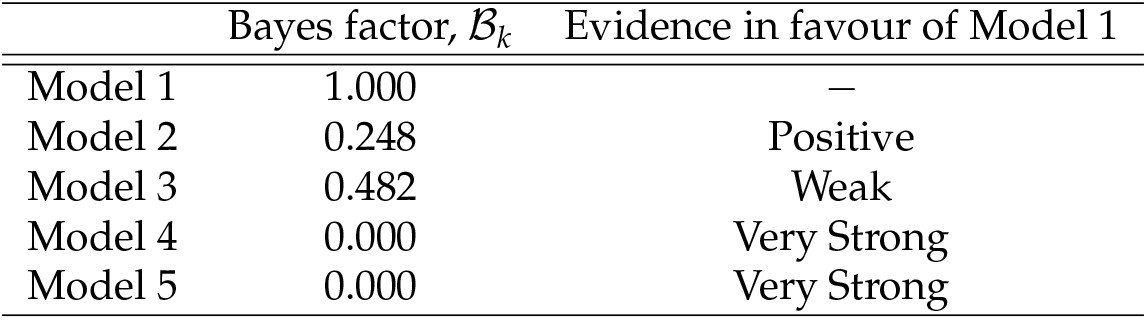
Bayes factor for each model, which describes the evidence in favour of Model 1 over Model *k*. A Bayes factor close to 1 indicates limited evidence in favour of Model 1 over Model *k*, and a Bayes factor close to 0 indicates very strong evidence in favour of Model 1 over Model *k* [27].

| | Bayes factor, $\mathcal{B}_k$ | Evidence in favour of Model 1 |
| --- | --- | --- |
| Model 1 | 1.000 | — |
| Model 2 | 0.248 | Positive |
| Model 3 | 0.482 | Weak |
| Model 4 | 0.000 | Very Strong |
| Model 5 | 0.000 | Very Strong |

**Figure 4:**
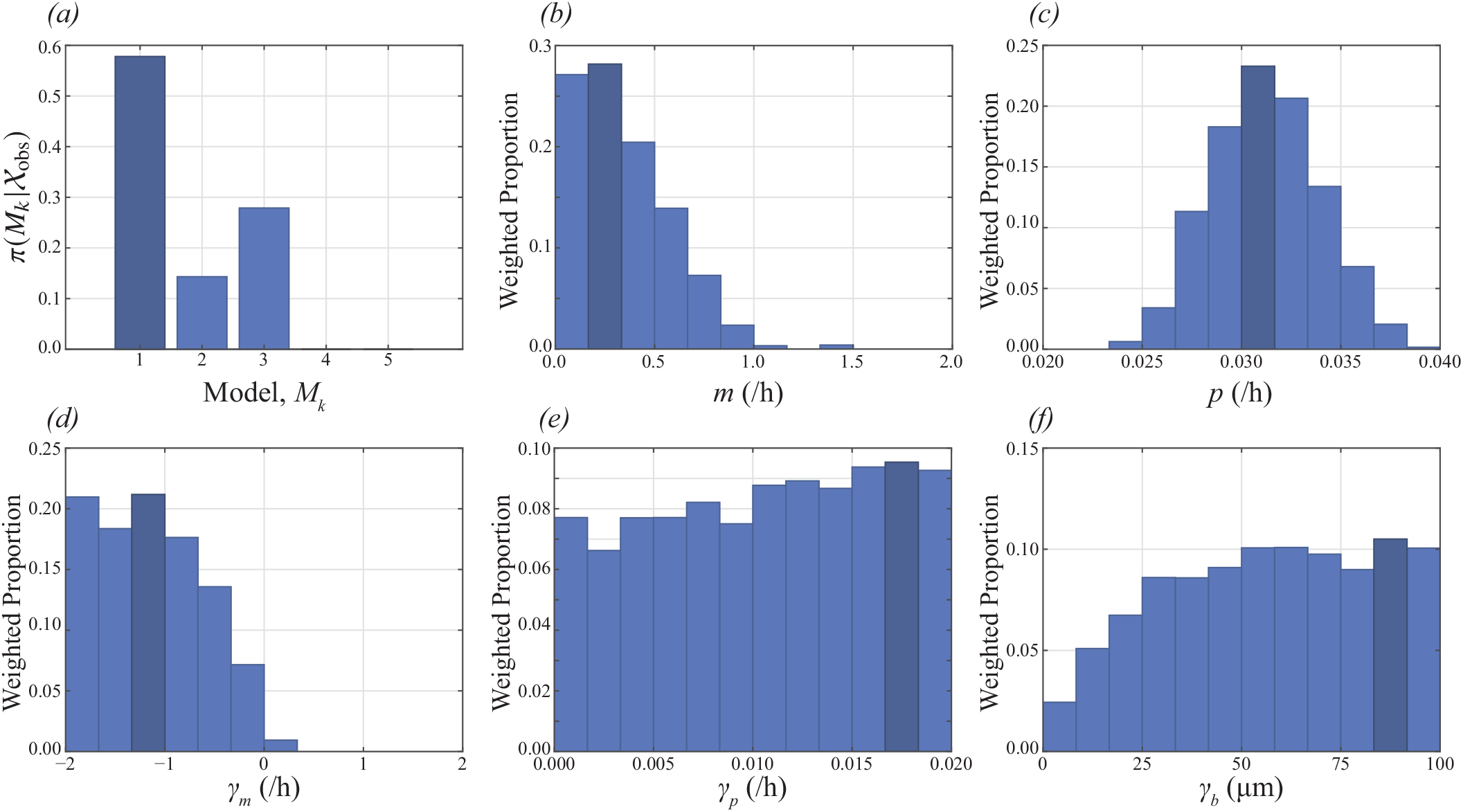
(a) Posterior for the model index, 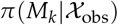, showing that Model 1 (the full model) is the posterior mode. (b)–(f) Marginal posterior distributions for each parameter in Model 1, shown as weighted histograms. In all cases, the posterior mode is indicated in dark blue.

We now focus on results for the full model (Model 1), which has the highest posterior density. In figure 4*b-f* we show marginal posterior distributions for each parameter in Model 1, and in figure 5 we compare the experimental data from four of the nine experiments to the calibrated model (in the supporting material, we show these results for all nine experiments). Overall, we find an excellent match between the model and experimental data, which has not been seen across a range of initial densities for this kind of experimental data. In addition to matching the density profile(figure 5*m–p*) and population (figure 5*u–x*), we find that the calibrated IBM is able to capture information about the spatial structure of cells, specifically, the pair correlation function (figure 5*q–t*). We perform a posterior predictive check for each summary statistic by producing 50% and 95% prediction intervals (PI) that characterise both the parameter uncertainty and stochasticity described by the model. The summary statistics produced from the experimental data almost always lie completely within the 95% PI, further indicating that the calibrated model is consistent with the experimental data across the range of initial densities. While we have not presented these results for Models 2 and 3, which have non-zero posterior density, the nature of ABC means that all accepted samples lie a similar distance to the experimental data.

**Figure 5:**
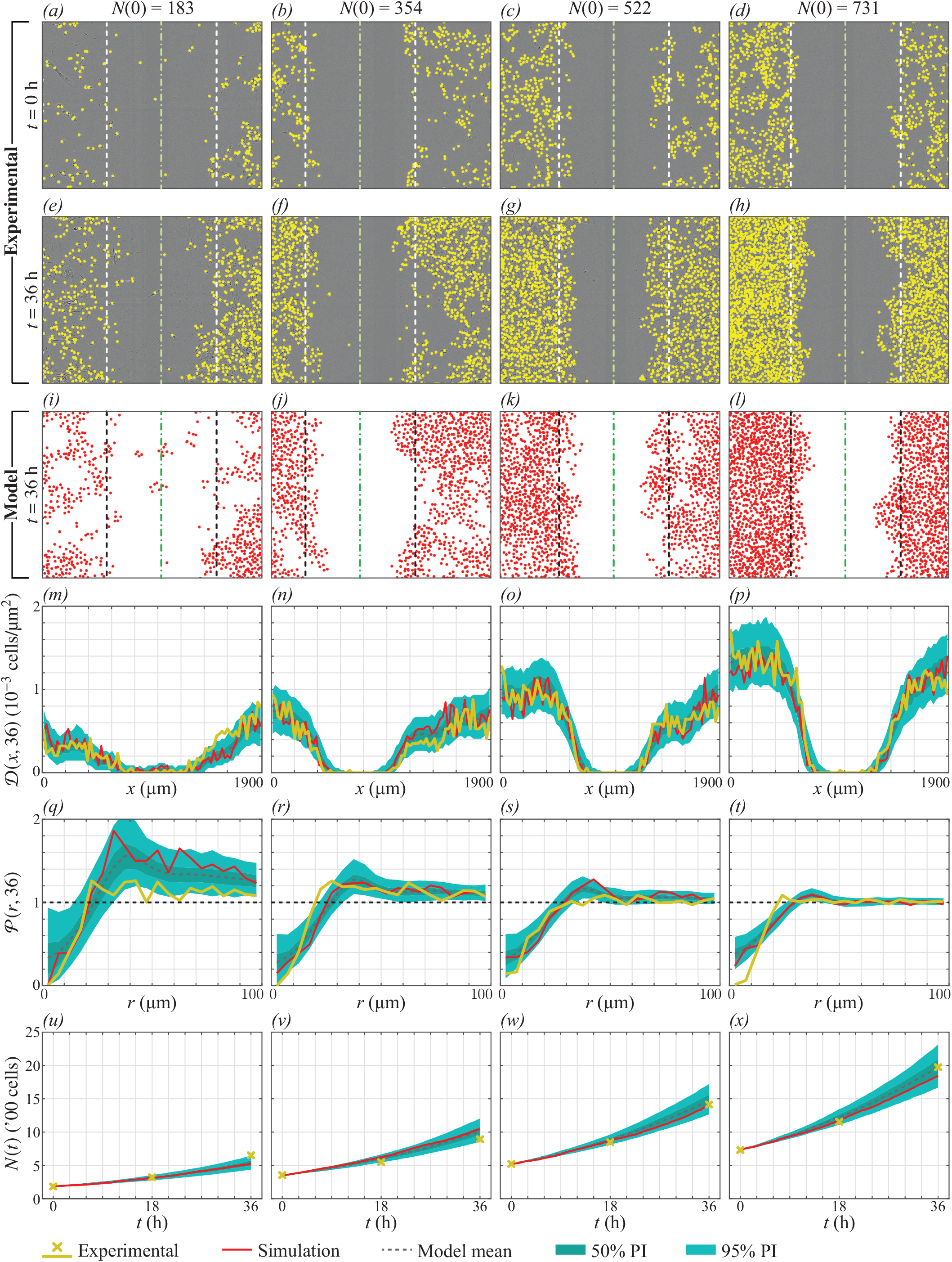
Comparison between experimental data and model estimate. Dashed lines in (a)–(l) give the scratched region that was used to compare the observed and simulated density profile, where green dash-dot lines give the approximate centre of the scratch at *t* = 0 h. Simulation results in (i)–(x) refer to a single realisation of the IBM using the parameter combination that gave the smallest discrepancy. In (m)–(x) the model mean, 50% prediction interval (PI) and 95% PI were produced using 500 realisations of the IBM using parameters resampled from the posterior distribution. Results for all nine experiments are provided supporting material (section 6).

Results in figure 4*d* suggest that *γ_m_ <* 0, so that crowding *increases* motility. This is consistent with mean-field models such as the porous Fisher equation [43] where the diffusivity increases with local density, but contrasts to other non-linear diffusion models where cell motility decreases with crowding [44]. This observation also explains why model realisations with small values of the motility rate, *m*, are able to match the data (this is seen in figure 4*b*), since a value *γ*_*m*_ < 0 allows motility in crowded regions if *m* ≪ 1. Interestingly, these results are less clear in the case where the pair correlation function is neglected (supporting material, section 2.3), which highlights the importance of considering spatial structure when studying these interactions. The increase of motility due to crowding may correspond to mechanical interactions, such as volume exclusion, in regions of very high cell density. It is trivial to add mechanisms to the IBM, and future work may examine *γ_m_* in the case volume exclusion [26], or other kinds of mechanical interactions [45–47], are included as additional mechanisms. Alternatively, the inclusion of non-monotonic interaction kernels [48] may allow movement to increase for agents close together, and decrease in crowded regions.

An interesting result is that the directional bias is included in the models with the highest posterior density (Models 1 and 3), but examining the marginal posterior for *γ*_*b*_ (figure 4*f*), we see that the strength of this bias may not be identifiable: the posterior distribution is relatively flat without a clear mode. These results might suggest that, past a certain point, increasing the strength of the directional bias has negligible effect. We verify these observations by widening the prior distribution for *γ*_*b*_ by a factor of two in the supporting material (section 2.2). To obtain more information about the strength of the directional bias, more detailed data, such as time-lapse cell tracking data, may be required [47].

Results in figure 4*e* indicate that the proliferation interaction strength parameter, *γ_p_*, appears to be unidentifiable [49], since the posterior distribution contains no well defined maxima. Figure 5*u–x* shows that population growth in both the experiments and calibrated model appears to be exponential, so we do not see crowding effects on proliferation in these experiments [28,50]. We verify this by performing model selection with three additional models (Models 6–8) that respectively correspond to Models 1– 3 with *γ*_*p*_ = 0 (supporting material, section 4). These additional results show that the distributions for Models 6–8 are similar to those for Models 1–3 and confirm that crowding effects on proliferation, such as contact inhibition, are simply not seen in these experiments. Early-time data often illustrates exponential growth for a variety of growth laws and experiments must be conducted over a longer period of time to identify the appropriate growth function [28].

## 4 Conclusion

The ability of common mean-field models, such as the Fisher-Kolmogorov equation and its generalisations, to match experimental data across a range of densities is rarely tested as typical experimental protocols do not vary the initial number of cells. These models typically assume either or both density-dependent proliferation and densityindependent motility [3, 7–12]. By modelling density-dependent interactions which affect motility, proliferation and directional bias, we calibrate a mathematical model that simultaneously describes scratch assay data across a range of densities. Using model selection, we quantitatively assess which interactions are most important. Our results provide an indication of how density affects the behaviour cells in our experiments. In opposition to common modelling assumptions [7], proliferation appears to be unaffected as density increases whereas cells become more motile as density increases. This information provides a hint about the kind of partial differential equation is most appropriate, such as

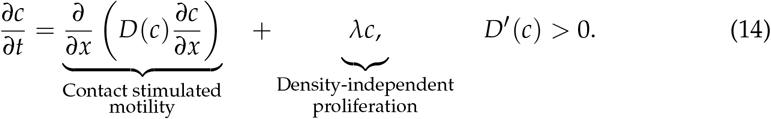

may be a more appropriate mean-field model for this kind of experimental data. Our findings agree with some areas of the literature [22–24], but contrast with studies that suggest density decreases motility [20, 21], or that motility is independent of density [3, 19]. Our finding that exponential growth describes the data is consistent with observations by Vittadello *et al.* [51], who point out that a loss of contact inhibition if a hallmark of cancer [52]. Despite this, the assumption of logistic growth is common in the mathematical modelling literature [32, 53, 54]. Our study demonstrates that exponential and logistic growth are not always distinguishable from typical experimental data [28]. Parameter identifiability [49] should be considered when calibrating logistic growth models to scratch assay data.

We study collective cell behaviour using a mathematical model which incorporates density-dependent interactions affecting proliferation, motility and directional bias. Applying SMC, which penalises models with high dimensionality of the unknown parameters, our study suggests the minimal model required to match the experimental data. Two of the primary advantages of our IBM approach is the ability to precisely replicate the experimental initial condition; and, the ease of which new mechanisms can be incorporated into, and removed from, the model. Our approach can, therefore, be applied to quantify experimental evidence for more complex mechanisms including chemotaxis [15, 55], mechanotaxis [56], and generalised growth laws [53], as well as comparing more complicated choices of interaction kernel [34]. Cell aspect ratio [23] can be incorporated into the model using asymmetric choices of interaction kernels, however would require more detailed experimental data, such as that provided by machine vision. We do not pursue such extensions here since we find that our simpler modelling framework already provides a good match to experimental data across a range of densities.

## Acknowledgements

M.J.S. is supported by the Australian Research Council, M.J.P. is partly supported by Te Puū naha Matatini, a New Zealand Centre of Research Excellence, and W.J. is supported by a QUT Vice Chancellor’s Research Fellowship. We thank David Warne for technical advice. Computational resources and services used in this work were provided by the HPC and Research Support Group, Queensland University of Technology, Brisbane, Australia. We thank the four anonymous referee for their helpful suggestions.

## Author contributions

A.P.B. performed the research and wrote the paper. A.P.B. and W.J. processed the experimental data. All authors provided feedback and gave approval for final publication.

## Data accessibility

Experimental data, and code used in this work, are available on GitHub at github.com/ap-browning/scratchIBM.

