## Supplementary Material Document for "Identifying density-dependent interactions in collective cell behaviour"

### Supplementary Material for "Identifying density-dependent interactions in collective cell behaviour"

### 1 Experimental Data

| Experiment | $N(0)$ | $N(18)$ | $N(36)$ | Fold change |
|---|---|---|---|---|
| 1 | 183 | 322 | 652 | 3.56 |
| 2 | 299 | 528 | 927 | 3.10 |
| 3 | 354 | 549 | 893 | 2.52 |
| 4 | 404 | 636 | 1121 | 2.77 |
| 5 | 427 | 657 | 1077 | 2.52 |
| 6 | 522 | 849 | 1414 | 2.71 |
| 7 | 677 | 1200 | 2013 | 2.97 |
| 8 | 692 | 1285 | 1853 | 2.67 |
| 9 | 731 | 1155 | 1974 | 2.70 |

**Table S1:** Summary of experimental data showing the cell count, $N(t)$, and fold change, $N(36)/N(0)$, for all nine experiments.

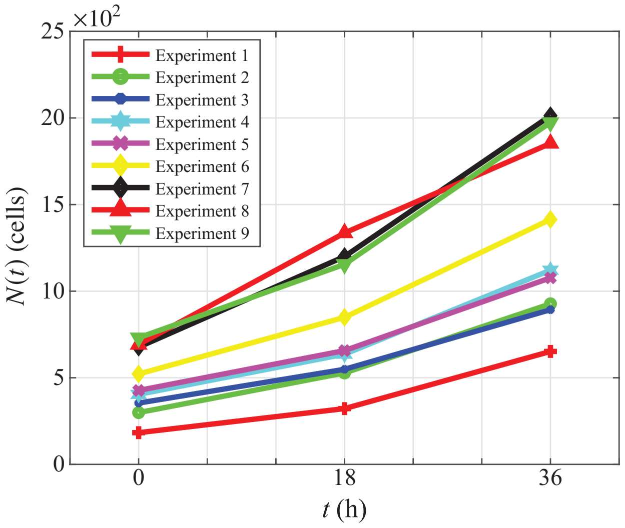

**Figure S1:** Summary of experimental data showing the cell count, $N(t)$, for all nine experiments.

#### 2 ABC algorithms

Here, we present the ABC rejection algorithm (algorithm 1) and ABC SMC algorithm (algorithm 2).

##### 2.1 ABC rejection algorithm

---

**Algorithm 1** ABC rejection sampling algorithm.

---

1: Draw parameter samples from the joint prior $\boldsymbol{\theta}_j \sim \pi(\boldsymbol{\theta})$.

2: Set discrepancy of $j$th sample $\kappa_j = 0$, experiment index $i = 1$.

  2.1: Set agent locations, $\{\mathbf{x}_n\}_{n=1}^{N(0)}$, to match experimental data $\mathbf{X}_{\text{obs}}^{(i)}$ at $t = 0$.

  2.2: Simulate model with parameters $\boldsymbol{\theta}_j$ for $t \leq 36$, storing the agent locations at $t = 18$ h and $t = 36$ h, denoted $\mathbf{X}_{\text{sim}}^{(i)}$.

  2.3: Update the discrepancy $\kappa_j \leftarrow \kappa_j + d(\mathbf{X}_{\text{obs}}^{(i)}, \mathbf{X}_{\text{sim}}^{(i)})$, where $d(\cdot, \cdot)$ is the discrepancy function.

  2.4: Move to the next replicate by setting $i = i + 1$ and repeat steps 2.1–2.4 until $i = 9$.

3: Repeat steps 1–2 until $10^5$ samples $\{\boldsymbol{\theta}_j, \varepsilon_j\}_{j=1}^{10^5}$ are simulated.

4: Order $\{\boldsymbol{\theta}_j, \kappa_j\}_{j=1}^{10^5}$ by $\kappa_j$ such that $\kappa_j < \kappa_{j+1}$.

5: Retain the first 1% ($\alpha = 0.01$) of prior samples $\boldsymbol{\theta}_j$, as posterior samples, $\{\boldsymbol{\theta}_j\}_{j=1}^{10^5\alpha}$.

---

#### 2.2 SMC algorithm

We apply the SMC model selection algorithm of Toni *et al.* [1], given in algorithm 2. We choose the perturbation kernel to be a multivariate Gaussian with independent components and variances approximately equal to the ABC rejection posterior variances [2].

---

**Algorithm 2** ABC SMC sampling algorithm for model selection with uniform priors [1].

1: Choose $\varepsilon_1, ..., \varepsilon_T$ such that $\varepsilon_k > \varepsilon_{k+1}$ and the desired total number of particles, $N_{\text{samples}}$. Set the population indicator $k = 1$.

2: Set the particle indicator $j = 1$.

- 2.1: Sample model indicator, $M_a^* \sim \pi(M_a)$.
- 2.2: If $k = 1$, sample proposal $\boldsymbol{\theta}^{**} \sim \pi_a(\boldsymbol{\theta})$ where $\pi_a(\boldsymbol{\theta})$ is the prior given model $M_a^*$. Go to step 2.4.
- 2.3: If $k > 1$, sample $\boldsymbol{\theta}^*$ from the subset of the previous population of particles for $M_a$, $\boldsymbol{\Theta}^{(a)}(k-1)$. If population is empty, return to 2.1. Perturb $\boldsymbol{\theta}^{**} \sim K(\boldsymbol{\theta}|\boldsymbol{\theta}^*)$, where $K(\cdot, \cdot)$ is a symmetric perturbation kernel. If $\pi_a(\boldsymbol{\theta}^{**}) = 0$, return to step 2.1.
- 2.4: Set discrepancy $\kappa = 0$, experiment index $i = 1$.
  - 2.4.1: Set agent locations, $\{\mathbf{x}_n\}_{n=1}^{N(0)}$, to match experimental data $\mathbf{X}_{\text{obs}}^{(i)}$ at $t = 0$.
  - 2.4.2: Simulate model $M_a$ with parameters $\boldsymbol{\theta}^{**}$ for $t \leq 36$, storing the agent locations at $t = 18$ h and $t = 36$ h, denoted $\mathcal{X}_{\text{sim}}^{(i)}$.
  - 2.4.3: Update the discrepancy $\kappa = \kappa + d(\mathbf{X}_{\text{obs}}^{(i)}, \mathbf{X}_{\text{sim}}^{(i)})$, where $d(\cdot, \cdot)$ is the discrepancy function.
  - 2.4.4: If $\kappa > \varepsilon_k$, reject particle and go back to 2.1. Else, move to next replicate by setting $i = i + 1$ and repeat steps 2.4.1–2.4.4 until $i = 9$.
- 2.5: Add $\boldsymbol{\theta}^{**}$ to the population of particles $\boldsymbol{\Theta}^{(a)}(k) = \{\boldsymbol{\theta}_j(k)\}_{j=1}^{N_{\text{samples}}^{(a)}}$, and calculate its weight as

$$w_j^{(a)}(k) = \begin{cases} 1, & k = 1, \\ \left( \sum_{j=1}^{N_{\text{samples}}^{(a)}} w_j^{(a)}(k-1) K(\boldsymbol{\theta}^{**}|\boldsymbol{\theta}_j(k-1)) \right)^{-1}, & k > 1. \end{cases}$$

- 2.6: Set $j = j + 1$ and repeat steps 2.1–2.6 until $j = N_{\text{samples}}$.

3. Set $k = k + 1$ and normalise the weights within each model. Repeat step 2 until $k = T$.

---

### 3 Pilot ABC results

#### 3.1 Pilot ABC to determine $\sigma$

In order to reduce the number of unknown parameters, we estimate and fix the kernel width parameter, $\sigma$. To do this, we perform a pilot run using ABC rejection, where $\pi(\sigma) = U(2, 30)$ using algorithm 1, the results of which are shown in figure S1. These results show a posterior mode of approximately $\sigma \approx 12\,\mu\text{m}$, which we treat as constant for the rest of this study. In this supporting material document, we also produce some results for $\sigma = 24\,\mu\text{m}$ to explore the sensitivity of this choice (figures S4, S6, S9 and S10).

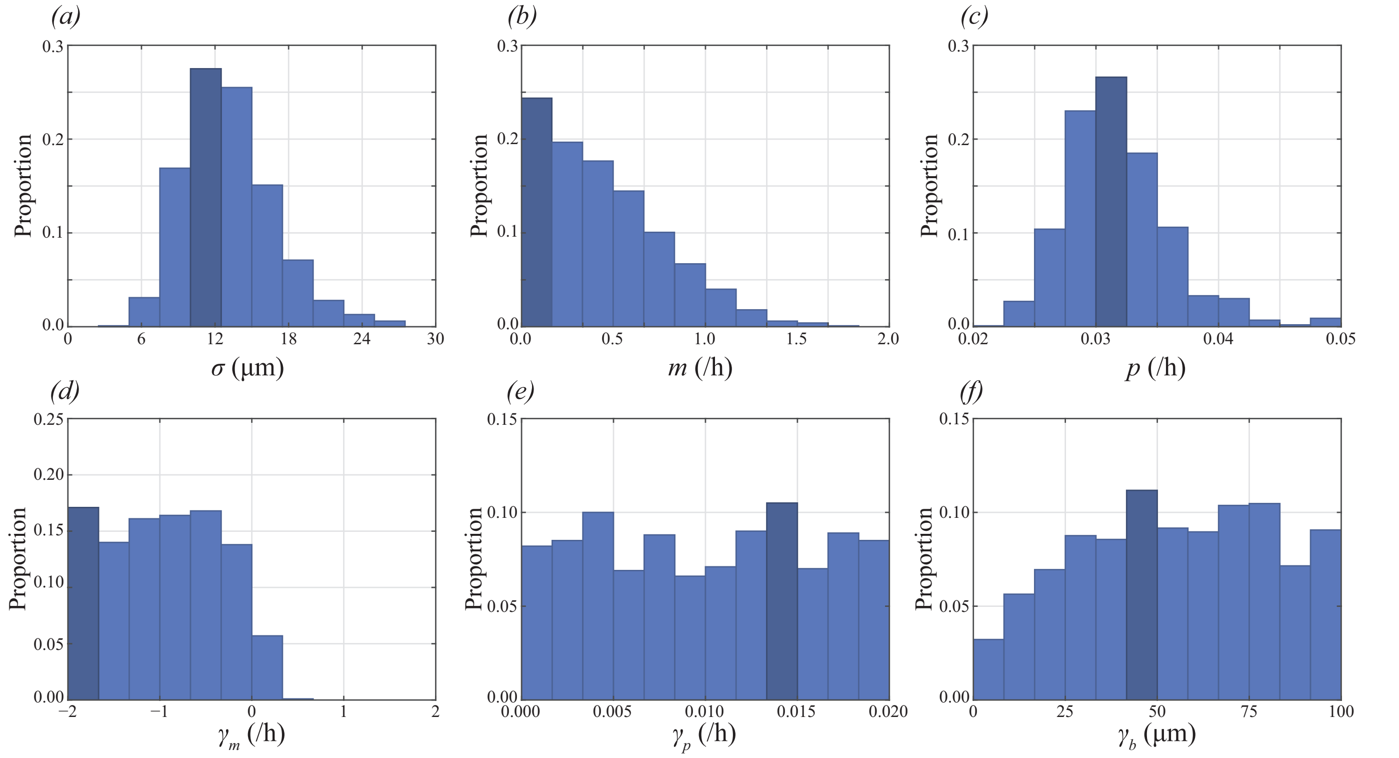

**Figure S1:** ABC rejection with $\sigma$ as an unknown with $\pi(\sigma) = U(2, 30)$.

#### 3.2 Wider priors

We notice in figure S1*e,f* that the posterior support appears to cover the prior support. To investigate this, we widen the corresponding priors for $\gamma_p$ and $\gamma_b$ by a factor of two and perform ABC rejection. The results are shown in figure S2. These results suggest that $\gamma_p$ and $\gamma_b$ are non-identifiable since the re-calculated posteriors are relatively flat and cover the same interval as the widened prior.

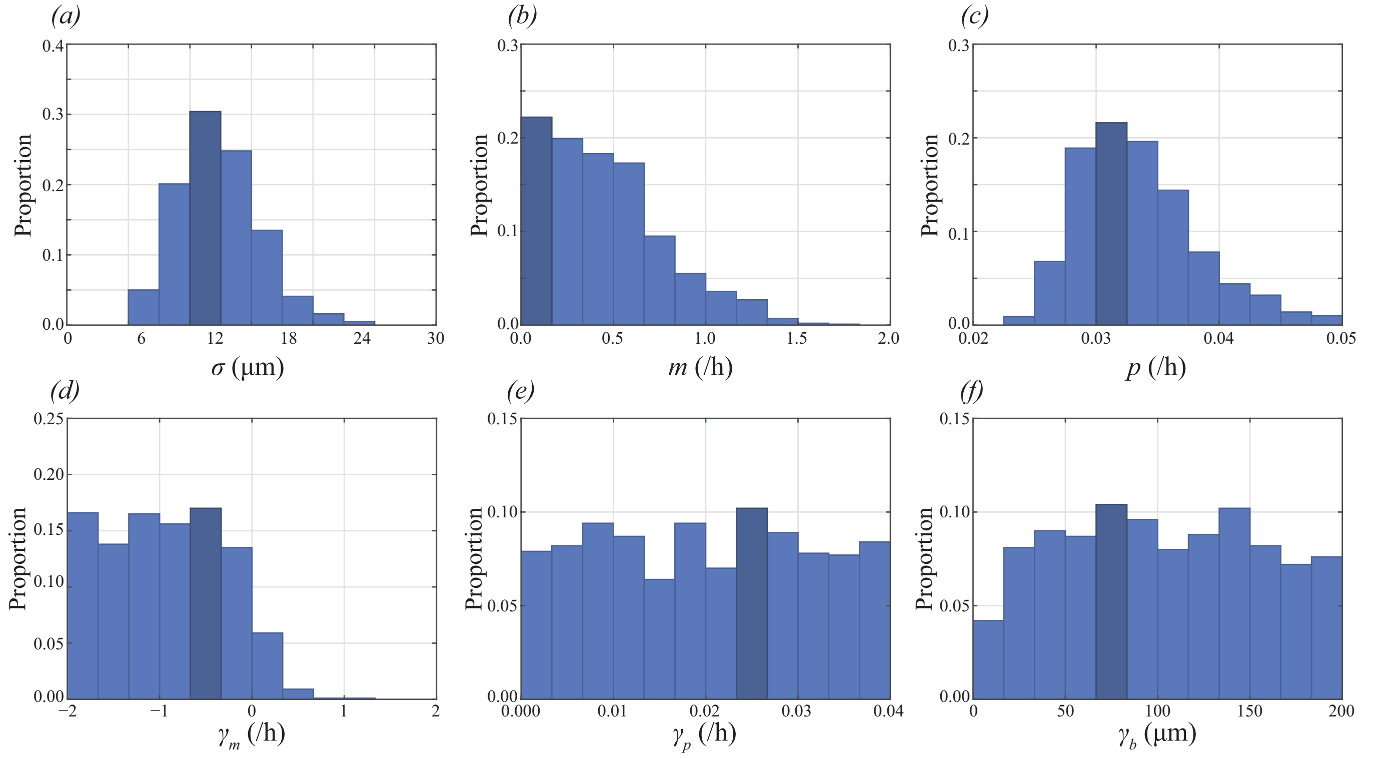

**Figure S2:** Figure S1 with wider priors for $\gamma_p$ and $\gamma_b$.

##### 3.3 Excluding $\mathcal{P}$ from inference

To investigate the amount of information contained in the pair correlation function, $\mathcal{P}$, we perform ABC rejection in the case $\mathcal{P}$ is removed from the distance metric so that

$$d(\mathbf{X}_{\text{obs}}, \mathbf{X}_{\text{sim}}) = \sum_{t \in \{18,30\}} \left( \frac{[N_{\text{sim}}(t) - N_{\text{obs}}(t)]^2}{N_{\text{obs}}(t)^2} + \frac{\sum_{j=I_{\text{mid}}-20}^{I_{\text{mid}}+20} [\mathcal{D}_{\text{sim}}(j,t) - \mathcal{D}_{\text{obs}}(j,t)]^2}{\sum_{j=I_{\text{mid}}-20}^{I_{\text{mid}}+20} \mathcal{D}_{\text{obs}}(j,t)^2} \right). \quad (1)$$

The results are shown in figure S3. We see a large reduction in information in the posteriors in figure S3 compared to figure S1. In particularly, the sign of $\gamma_m$ is less clear in the case the pair correlation is excluded.

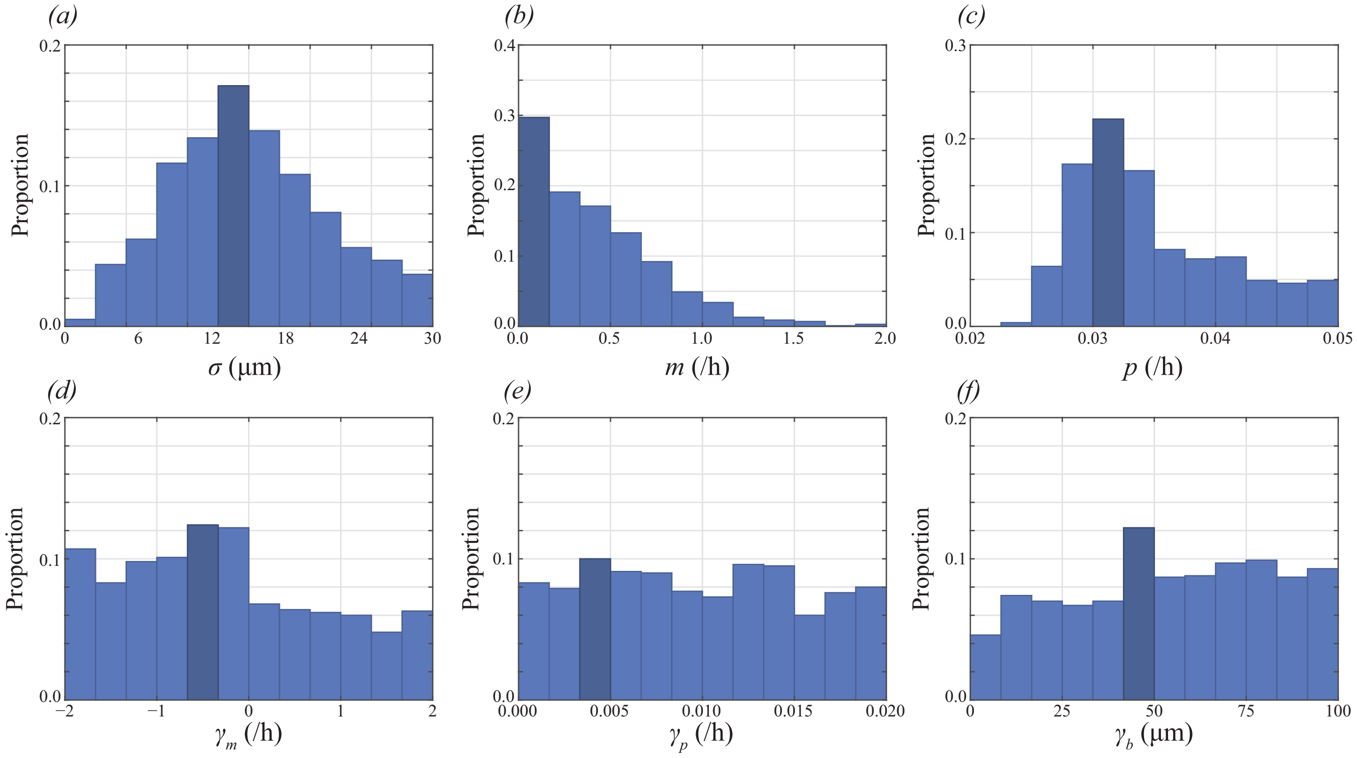

**Figure S3:** Figure S1 where summary statistics relating the the pair correlation, $\mathcal{P}$ are excluded.

### 4 Quantile plots

To determine the appropriate ABC SMC sequence of thresholds, we produce a quantile plot of the distance metric obtained from 100,000 prior samples where $\sigma = 12\,\mu\text{m}$ (figure S4*a*) and $\sigma = 24\,\mu\text{m}$ (figure S4*b*). In each case, we choose the final discrepancy, $\varepsilon_U$, to correspond to an ABC rejection rate of approximately 1%. We choose the sequence base upon acceptance probabilities of approximately 50%, 25%, 12.5%, 6.25%, 3.125%, 1.5625% and 1% [2]. The sequence of thresholds for results in the main document where $\sigma = 12\,\mu\text{m}$ is $\{9.6, 7.3, 6.3, 5.5, 4.9, 4.6, 4.4\}$; and the sequence of thresholds for results in this supporting material document where $\sigma = 24\,\mu\text{m}$ is $\{11.8, 9.7, 8.4, 7.5, 6.8, 6.3, 6.0\}$.

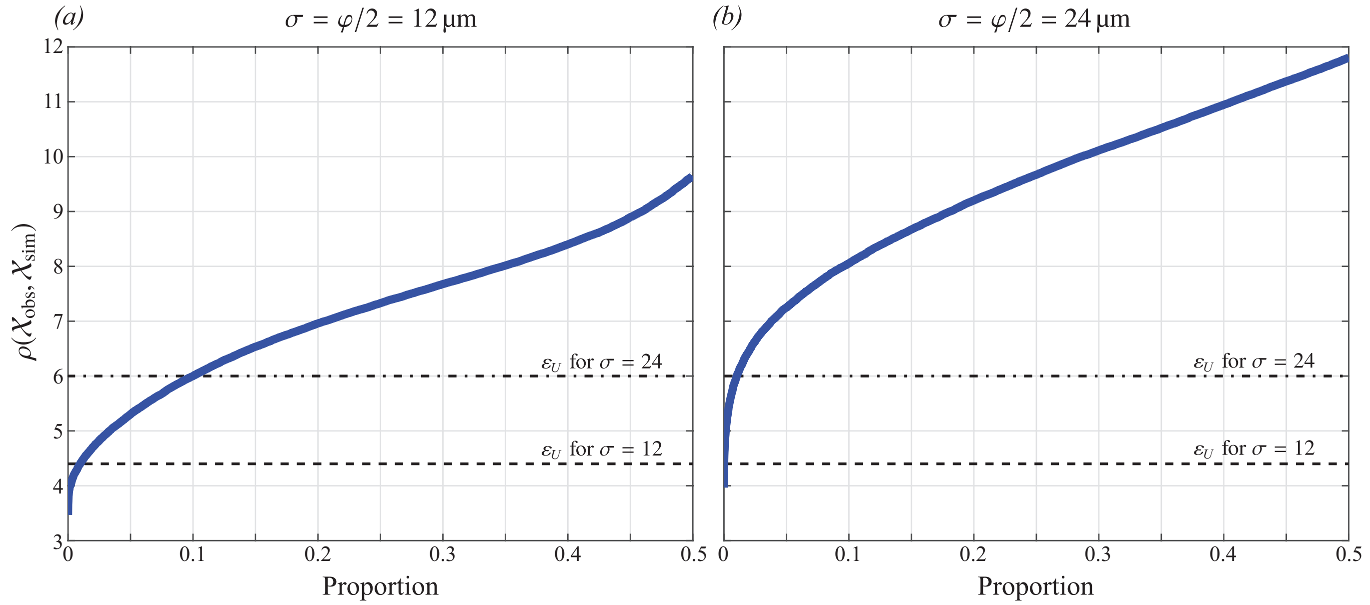

**Figure S4:** Quantile plot of the distance metric from 100,000 prior samples with (a) $\sigma = 12\,\mu\text{m}$; and, (b) $\sigma = 12\,\mu\text{m}$. The final discrepancy, $\varepsilon_U$, is indicated by a horizontal line in each case.

### 5 Exponential growth

To explore the possibility that $\gamma_p = 0$, which corresponds to exponential growth, we consider three additional models: Models 6, 7 and 8, that correspond to Models 1, 2 and 3, with $\gamma_p = 0$.

| | $\boldsymbol{\theta}_k$ | Density Dependence |
|---|---|---|
| Model 1 | $(m, p, \gamma_m, \gamma_p, \gamma_b)$ | Proliferation, Motility and Direction |
| Model 2 | $(m, p, \gamma_p, \gamma_b)$ | Proliferation and Direction |
| Model 3 | $(m, p, \gamma_m, \gamma_p)$ | Proliferation and Motility |
| Model 4 | $(m, p, \gamma_p)$ | Proliferation only (Fisher-Kolmogorov [3,4]) |
| Model 5 | $(m, p)$ | None (Skellam [5]) |
| Model 6 | $(m, p, \gamma_m, \gamma_b)$ | Motility and Direction |
| Model 7 | $(m, p, \gamma_b)$ | Direction only |
| Model 8 | $(m, p, \gamma_m)$ | Motility only |

**Table S2:** Here we consider three additional models: Models 6, 7 and 8 correspond to Models 1, 2 and 3, where $\gamma_p = 0$.

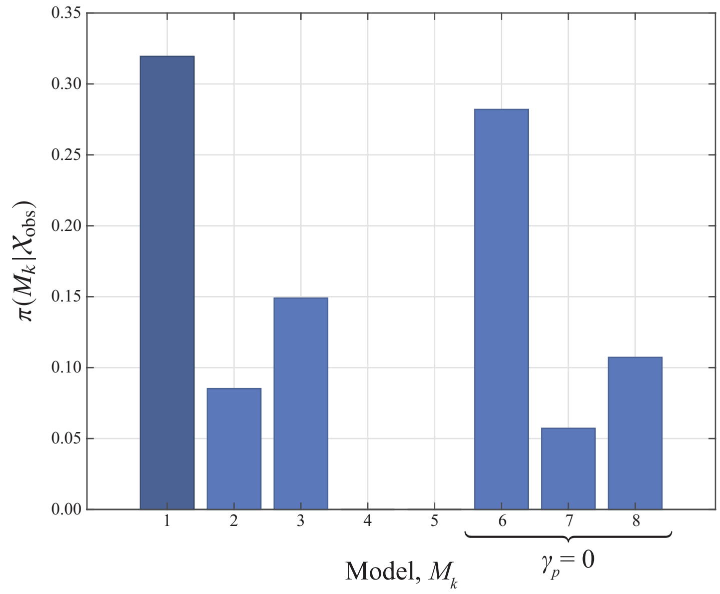

**Figure S5:** Figure 4 of the main document where we consider three additional models: Models 6, 7 and 8 correspond to Models 1, 2 and 3 where we set $\gamma_p = 0$ to remove the proliferation interaction from the model.

| | Bayes factor, $\mathcal{B}_k$ | Evidence in favour of Model 1 |
|---|---|---|
| Model 1 | 1.000 | – |
| Model 2 | 0.267 | Positive |
| Model 3 | 0.467 | Weak |
| Model 4 | 0.000 | Very Strong |
| Model 5 | 0.000 | Very Strong |
| Model 6 | 0.883 | Weak |
| Model 7 | 0.179 | Positive |
| Model 8 | 0.336 | Weak |

**Table S3:** Bayes factor for each model, which describes the evidence in favour of Model 1 over Model $k$. A Bayes factor close to 1 indicates limited evidence in favour of Model 1 over Model $k$, and a Bayes factor close to 0 indicates very strong evidence in favour of Model 1 over Model $k$ [1]. Here, Models 6, 7 and 8 correspond to Models 1, 2 and 3, where $\gamma_p = 0$.

### 6 Model selection for $\sigma = 24\,\mu\text{m}$

Here, we reproduce results from figure 5 of the main document, in the case we fix $\sigma = 24\,\mu\text{m}$.

Results in figure S6*a* differ from those in the main document, in that Model 2 now has the highest posterior density. We show the marginal distributions for each parameter in Model 2 in figure S6*b*–*f*. These results are consistent with the main document in showing that models without a density dependent motility mechanism (Models 4 and 5) are unable to simultaneously match data from all nine experiments. Examining results in figure S10 shows that $\sigma = 24\,\mu\text{m}$ is not able to match the spatial structure in the experimental data as closely as $\sigma = 24\,\mu\text{m}$, results for which are shown in figure S8.

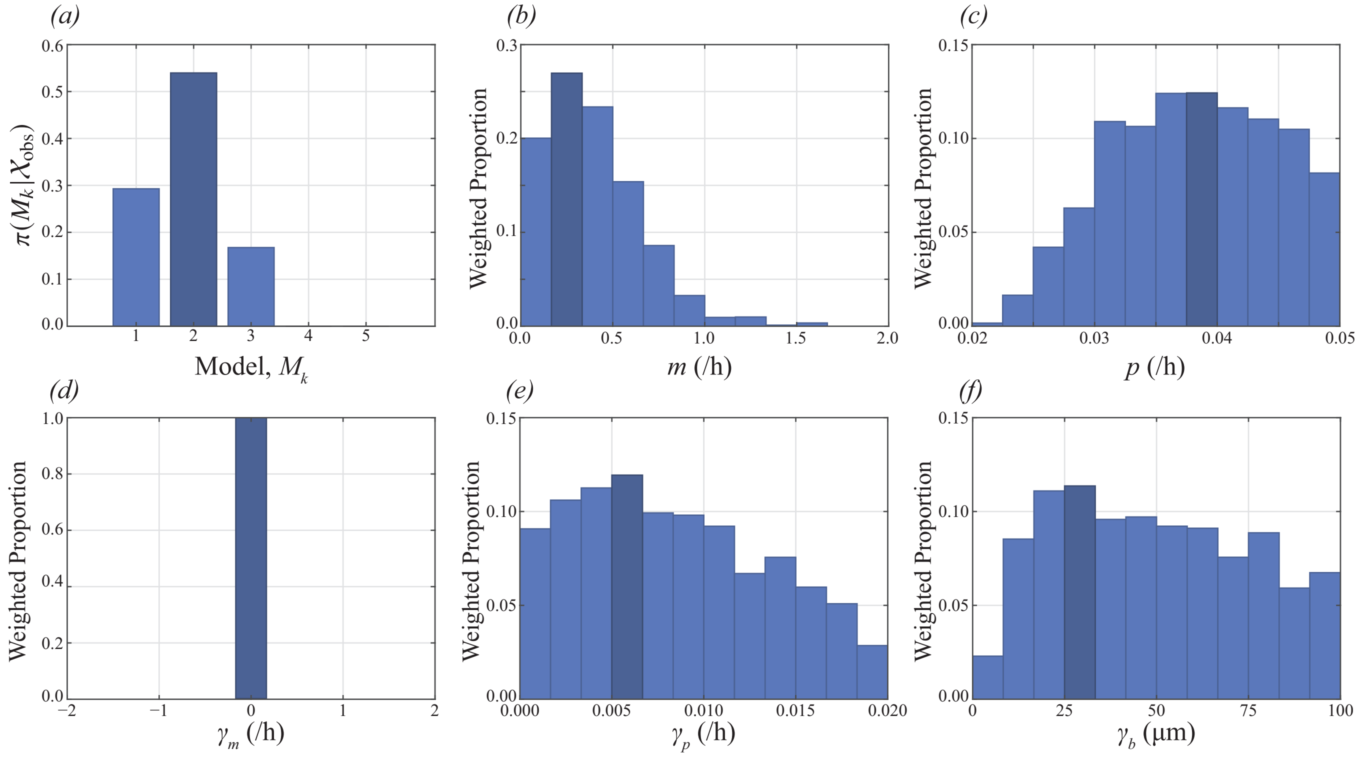

**Figure S6:** Reproduction of results in figure 6 of the main document, with $\sigma = \varphi = 24\,\mu\text{m}$. (a) Posterior for the model index, $\pi(M_k|\mathcal{X}_{\text{obs}})$, showing that Model 2 (density-independent motility) as the posterior mode. (b)–(f) Marginal posterior distributions for each parameter in Model 2, shown as weighted histograms.

### 7 Results for all nine experiments

In the main document, we show results for experimental replicates 1, 3, 6 and 9 at $t = 36\,\text{h}$ in figure 6. Here, we reproduce figure 6 and show results for all nine experimental replicates at both $t = 18\,\text{h}$ and $t = 36\,\text{h}$.

In section 7.1 we show these results for $\sigma = 12\,\mu\text{m}$, the value from the main document. In section 7.2 we show these results for $\sigma = 24\,\mu\text{m}$.

##### 7.1 Full results for $\sigma = 12\,\mu\text{m}$

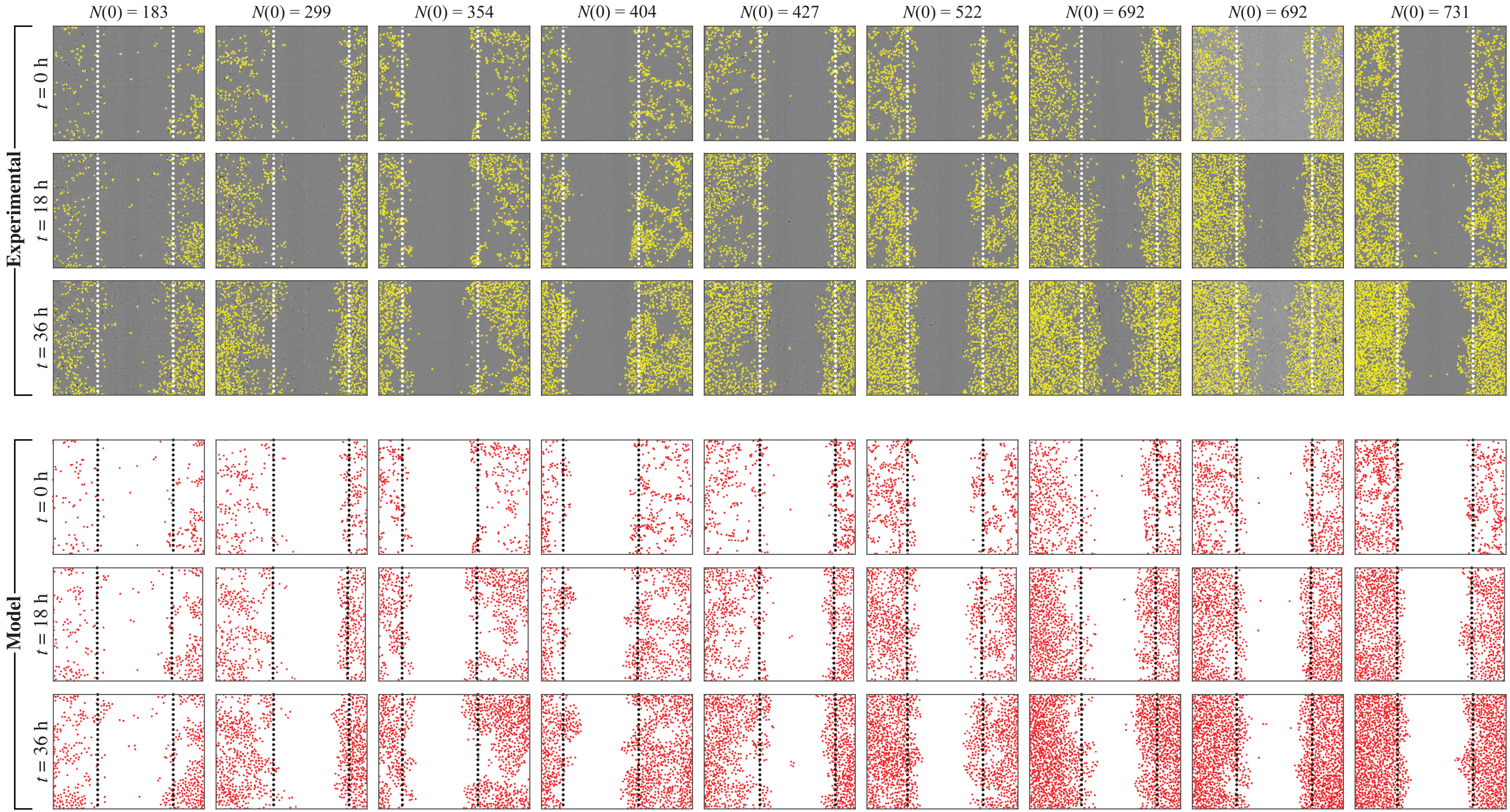

**Figure S7:** Reproduction of results in figure 6*a*–*l* of the main document, for all replicates, for $\sigma = 12\,\mu\text{m}$.

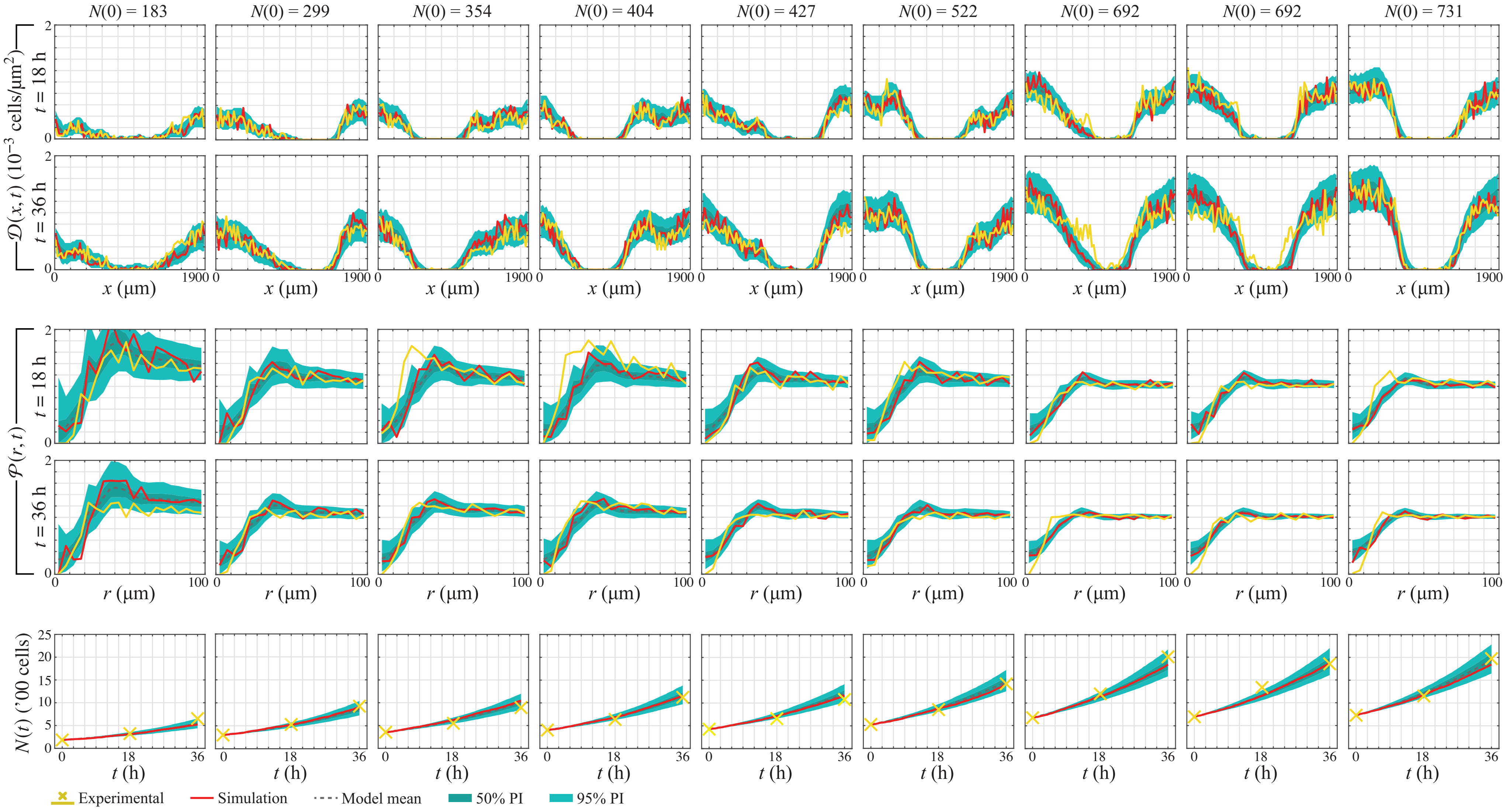

**Figure S8:** Reproduction of results in figure 6*m*–*x* of the main document, for all replicates, for $\sigma = 12\,\mu\text{m}$.

#### 7.2 Full results for $\sigma = 24\,\mu\text{m}$

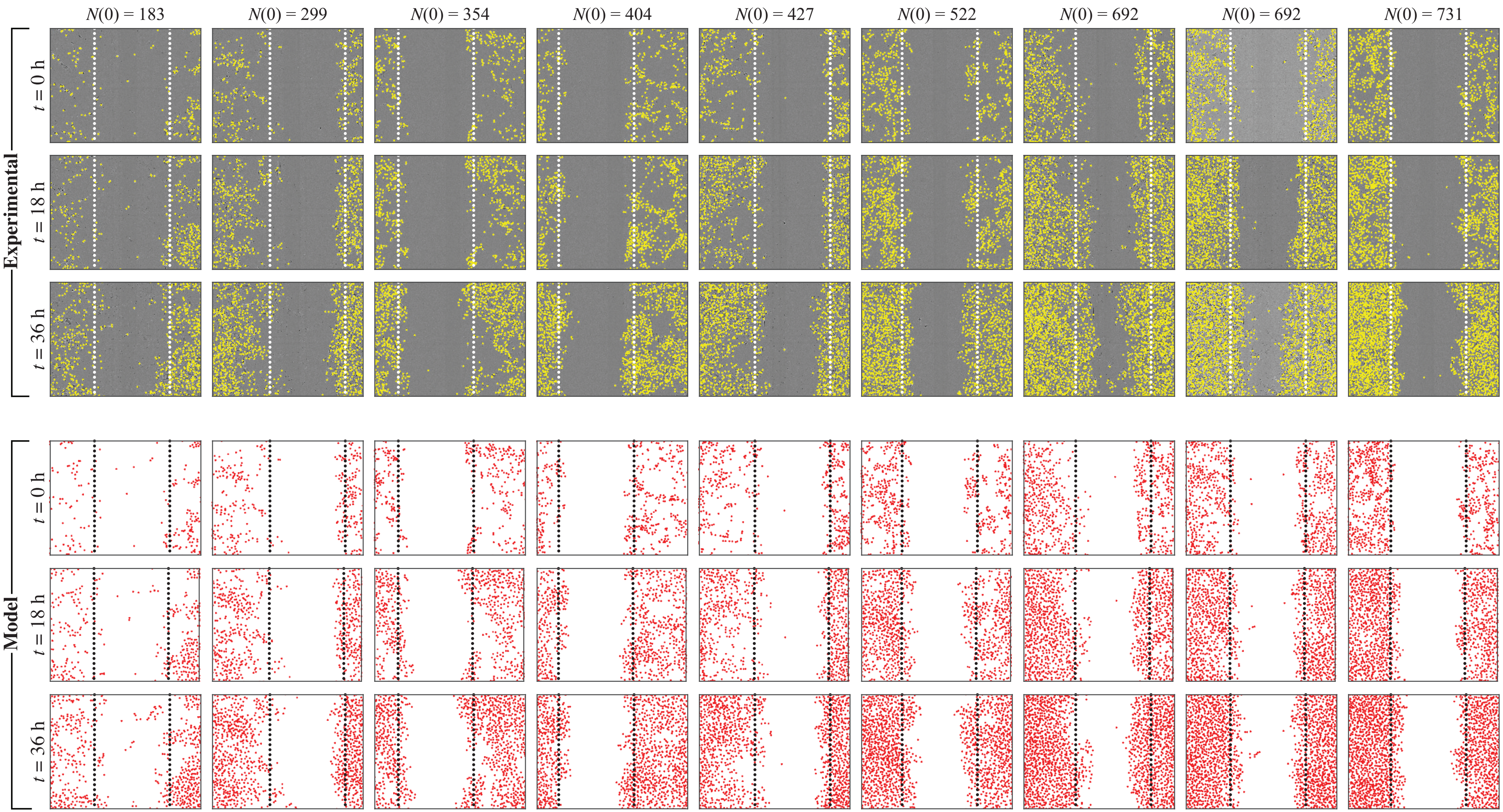

**Figure S9:** Reproduction of results in figure 6*a*–*l* of the main document, for all replicates, for $\sigma = 24\,\mu\text{m}$.

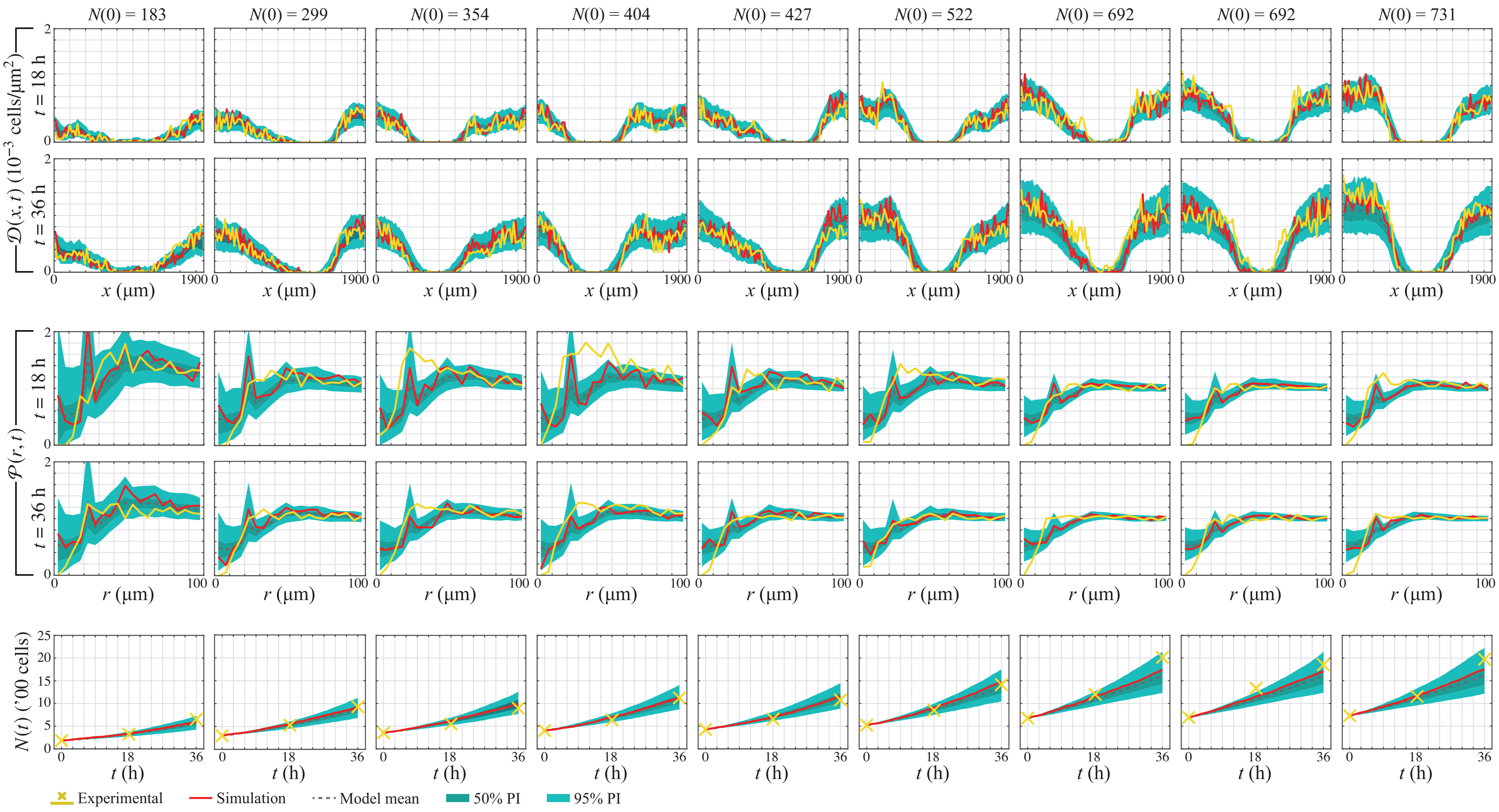

**Figure S10:** Reproduction of results in figure 6*m*–*x* of the main document, for all replicates, for $\sigma = 24\,\mu\text{m}$.
